# Multisensory integration (MSI) by polymodal sensory neurons dictates larval settlement in a brainless cnidarian larva

**DOI:** 10.1101/2022.11.02.514770

**Authors:** Sydney Birch, David Plachetzki

## Abstract

Multisensory integration (MSI) combines information from more than one sensory modality to elicit behaviors distinct from unisensory behaviors. MSI is best understood in animals with complex brains and specialized centers for parsing sensory information, but the dispersive larvae of sessile marine invertebrates utilize multimodal environmental sensory stimuli to base irreversible settlement decisions on, and most lack complex brains. Here, we examined the sensory determinants of settlement in actinula larvae of the hydrozoan *Ectopleura crocea* (Cnidaria), which possess a diffuse nerve net. A factorial settlement study revealed that photo-, chemo-, and mechano-sensory cues each influence the settlement response, which was complex and dependent on specific combinations of cues, therefore indicating MSI. Mechanosensory cues either inhibited or enhanced settlement rates depending on the presence or absence of chemical and light cues in the environment. Sensory gene expression over development peaked with developmental competence to settle, which in actinulae, requires cnidocyte discharge. Transcriptome analyses also highlighted several deep homological links between cnidarian and bilaterian mechano- chemo- and photo-sensory pathways. Fluorescent *in situ* hybridization studies of candidate transcripts suggested cellular partitioning of sensory function among the few cell types that comprise the actinula nervous system, where ubiquitous polymodal sensory neurons with putative chemo- and photo-sensitivity interface with mechanoreceptive cnidocytes. We propose that a simple multisensory processing circuit, involving polymodal chemo/photosensory neurons and mechanoreceptive cnidocytes, is sufficient to explain MSI in actinulae settlement. Our study demonstrates that MSI is not exclusive to complex brains, but likely predated and contextualized their evolution.

## Introduction

A distinguishing feature of animals is their exquisite capacity to receive sensory information from the environment and integrate it into behavior. Animals may integrate sensory signals from individual modalities (e.g. vision or taste), or they may perform multisensory integration (MSI), which combines information from more than one modality to elicit behaviors that are distinct from those elicited by unisensory behaviors (Stein et al. 1989, 2009, 2014; Stein and Meredith 1993; Alvarado et al. 2007; Otto et al. 2013; Stevenson et al. 2014).

MSI is best understood in bilaterian animals with complex nervous systems that include specialized centers for information processing and exchange (Stein 1998; Otto et al. 2013; Stein et al. 2014; Ghosh et al. 2017; Currier and Nagel 2020). Classically, MSI studies are conducted at the neuron level where neuronal signals are processed in the brain to make behavioral decisions (Meredith and Stein 1983; Stein 1998; Stein and Stanford 2008; Otto et al. 2013). However, much less research has been conducted on organisms without complex nervous systems or brains. Moreover, it has been shown that zoospores of an *Allomyces* fungus utilize a multisensory system involving chemo- and photo-taxis (Swafford and Oakley 2018) which suggests the possibility that MSI evolved prior to the evolution of complex nervous systems.

Sensory integration is critically important for sessile marine invertebrates that utilize larvae for dispersal and often make irreversible settlement decisions. Because some sensory cues may be better indicators of site quality than others, it has been proposed that larvae place emphasis on select cues, leading to a hierarchy of sensory cues that determine where settlement occurs (Kingsford et al. 2002; Müller and Leitz 2002; Woodson et al. 2007; Hodin et al. 2018). However, MSI has yet to be demonstrated in marine invertebrate larval settlement, and little is known about the potential for MSI in such organisms that lack complex nervous systems or brains.

The marine hydrozoan *Ectopleura crocea* is a benthic colonial species with a pan-global distribution in temperate coastal regions. Unlike many other cnidarians, *E. crocea* possesses an actinula larva (Fig 1C). Actinulae are motile lecithotrophic larvae that develop in the following five stages: the star embryo, preactinula, actinula, nematocyte-printing (settling) actinula, and settled actinula. *E. crocea* actinulae larvae begin settlement using a larval behavior called nematocyte-printing, which serves to tether actinula to the substrate (Yamashita 2003). However, little is known about the sensory cues that determine this process, the cell types that receive such information, or the underlying genetic machinery of sensation that coordinates such settlement decisions.

**Figure 1.**
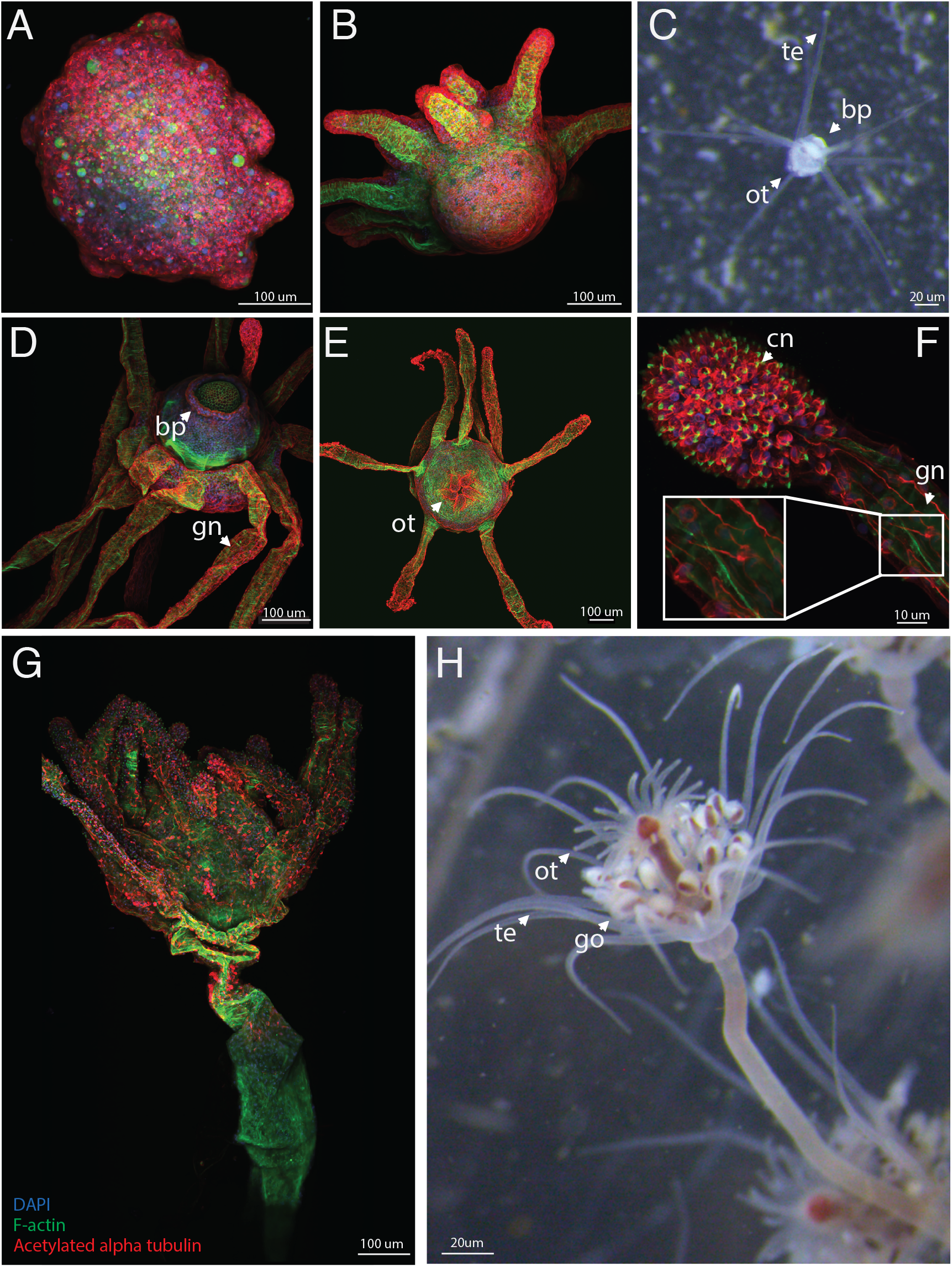
Nervous system development of *Ectopleura crocea* actinulae larvae. **A**,**B**,**D-G** Immunohistochemistry (IHC) staining of four developmental stages of *E. crocea* larvae where red staining corresponds to acetylated alpha-tubulin expression in neurons, green staining corresponds to contractile F-actin in muscles and stereocilia, and blue staining corresponds to DNA in nuclei. (**A**) Stage 1, star embryo. (**B**) Stage 2, preactinula. (**C**) Light micrograph of an actinula larva. The arrows point to the basal protrusion (bp) which attaches to the substrate during settlement, the tentacles (te), and the developing oral tentacles (ot). (**D**) The aboral end of a Stage 3-4 actinula larva. The arrows point to the basal protrusion (bp) and ganglion neurons (gn) in the tentacles. (**E**) Oral end of an actinula larva at stage 3-4. The arrow is pointing to the developing oral tentacles (ot). (**F**) High magnification of a stage 3-4 actinula tentacle. An abundance of cnidocytes (cn) are found at the tips of the tentacles. Additionally, ganglion neurons (gn) extend down the tentacles and connect to the actinular nerve net. (**G**) A metamorphosed juvenile polyp. (**H**) A light micrograph of an adult *E. crocea* polyp. The arrows point to the oral tentacles (ot), the aboral tentacles (te), and the gonophores (go).

Here we describe integrative studies on the sensory biology of settlement in the actinulae of *Ectopleura crocea*. Immunohistochemical studies of larval neural network development indicate that *E. crocea* larvae first possess a defined nervous system complete with robust tentacular cnidocytes and sensory neurons by the actinula stage of development. Next, to identify the sensory cues involved in settlement, we performed a factorial larval settlement study that investigated the effects of individual and combined cues (e.g., light, chemical (biofilm), and mechanical/surface texture) corresponding to three prominent sensory modalities. We found strong evidence of MSI during larval settlement where the highest rates of settlement occurred in the presence of all three sensory cues and where the effects of cues changed in the presence or absence of other cues, resulting in a sensory cue hierarchy. Developmental transcriptome analyses revealed previously undescribed, deep homological links with bilaterian sensory system development and a peak expression of sensory transduction components for each of the three modalities in actinulae stage larvae. Lastly, RNA Fluorescent *in situ* Hybridization (FISH) studies localize several prominent sensory transcripts including opsin (photosensitivity), PKD2L1, and PKD1L3 (chemosensitivity), and ASIC, Piezo, and TRPA (mechanosensitivity) to cnidocytes and their attendant sensory neurons in competent actinulae. Our results demonstrate MSI in the brainless actinula of *E. crocea* and suggest that this capacity is facilitated by the activities of polymodal, chemo-photo-mechanoreceptive sensory neurons with distant ancestry to unimodal primary sensory neurons known from bilaterian animals.

## Results

### Larval nervous systems reach full development by the actinula stage of development

The actinula stage of *E. crocea* possesses morphological and cellular features such as the basal protrusion and tentacles replete with cnidocytes, that are associated with larval settlement (Yamashita 2003). However, the structure of the nervous system throughout larval development has not been examined. We examined larvae at four developmental stages, including the metamorphosed juvenile polyp stage, using immunohistochemistry (IHC) and confocal microscopy (Fig 1). The state of the nervous system at the earliest stage, the star embryo (Fig 1A), appears granular and undifferentiated, which we interpret as a contiguous assemblage of neural progenitor cells (Rentzsch et al. 2017). The larval nervous system becomes increasingly differentiated as development proceeds from the preactinula (Fig 1B) to the actinula stage (Fig 1C-F), whereupon the tentacles are loaded with neurons and cnidocytes (Fig 1F). Additionally, the basal protrusion, the structure that contacts the substrate during settlement, contains a concentrated ring of neural cells (Fig 1D). These data are consistent with the actinula stage being the competent stage for settlement and suggest that actinulae have the capacity to integrate sensory information using the tentacles and the basal protrusion.

### Differences in the sensory environment impact larval settlement

Next, we investigated the sensory information actinula integrate in support of the settlement decision. We performed a factorial settlement study (Fig 2) assessing the impact of photosensory, chemosensory, and mechanosensory cues on larval settlement, which we analyzed using a three-way analysis of variance (ANOVA). Our experiment included 10 blocks (replicates) where each block had 28 treatments of all possible combinations of cues and examined a total of 2,800 individual actinula larvae (Fig 2A). While the blocking variable, which was the experimental day, did not influence the response, larval settlement did increase as the season progressed, similar to Yamashita et al (2003) (Fig S2). Each individual sensory cues significantly impacted settlement when assessed individually (light, *p*=0.0482; chemical, *p*=<2e-16; mechanical, *p*=<2e-16; Fig 2B-D; Table S1). Specifically, the rate of larval settlement increased significantly in the presence of light compared to darkness (*p*=0.0482; Fig 2B), and in the presence of the chemical cue compared to its absence (*p*=<2e-16; Fig 2C) but decreased in the presence of a mechanical cue (*p*=<2e-16; Fig 2D), when no other cues were present.

**Figure 2.**
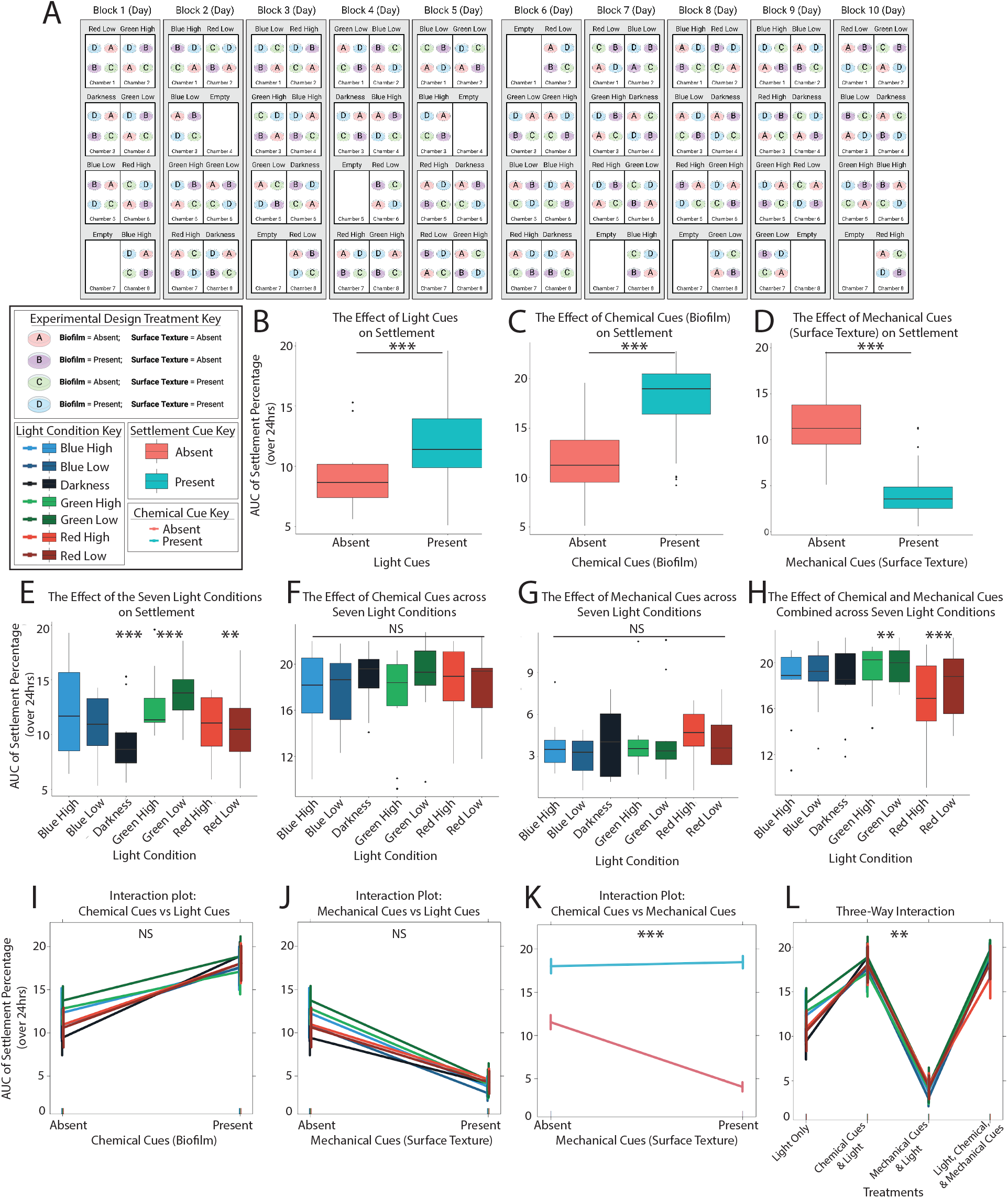
Settlement responses of actinula larvae to three sensory cues. (**A**) Cartoon depiction of the larval settlement experimental design: 7×2×2 Split-plot Randomized Complete Block Design (RCBD) with 10 replicates. The block represents an experimental day which is also a replicate. We included all four combinations of presence and absence of biofilm and surface texture which are depicted as A-D treatments, where each biofilm and texture combination was exposed to all 7 light conditions in a single replicate. (**B**) The individual effect of light cues on settlement. Larval settlement is significantly higher (*p*=0.0027) in the presence of a light cue compared to the absence of a light cue (darkness). (**C**) The individual effect of chemical cues (biofilm) on settlement. Larval settlement is significantly higher (*p*=<2e-16) in the presence of a chemical cue compared to the absence of one. (**D**) The individual effect of mechanical cues (surface texture) on settlement. Larval settlement is significantly lower (*p*=<2e-16) in the presence of a mechanical cue compared to the absence of one. (**E**) The effect of the seven light conditions on settlement. Larval settlement was significantly lower in darkness (*p*=0.00272) and in red wavelengths of light (*p*=0.01137) and significantly higher in green wavelengths of light(*p*=0.00167). (**F**) The effect of chemical cues (biofilm) across the seven light conditions. There were no significant differences in settlement in any wavelength of light (blue *p*=0.51; green *p*=0.60; red *p*=0.88) in the presence of a chemical cue. (**G**) The effect of mechanical cues (surface texture) across the seven light conditions. There were no significant differences in settlement in any wavelength of light in the presence of a mechanical cue (blue *p*=0.99; green *p*=0.79; red *p*=0.79). (**H**) The effect of chemical and mechanical cues combined across the seven light conditions. There was a significant increase in settlement in the presence of green wavelengths of light (*p*=0.02) and significantly lower settlement in red wavelengths of light (*p*=0.003). (**I**) The interaction of chemical cues and light cues. There was no significant interaction between these two cues (*p*=0.33). (**J**) The interaction between mechanical cues and light cues. There was no significant interaction between these two cues (*p*= 0.89). (**K**) The interaction between chemical and mechanical cues. There was a significant interaction between these two cues (*p*=<2e-16) with an additive effect in the presence of both cues. (**L**) The three-way interaction of chemical, mechanical, and light cues. Treatment A is the light cues only treatment, treatment B is light and chemical cues, treatment C is light and mechanical cues, and treatment D is all three cues. There is a significant three-way interaction (*p*=0.02).

The effect of photosensory cues on settlement (wavelengths of light) was investigated in the light-only treatment (Fig 2E) using contrast analyses, since this treatment included additional dimensions compared to chemical and mechanical cues (e.g., presence/absence of light, three wavelengths, and two intensities, totaling 7 light conditions; Fig 2A). Settlement increased in the presence of light compared to darkness (*p*=0.0027; Fig 2E and Fig 2B, Table S3). In addition, significantly higher settlement occurred in green wavelengths of light (*p*=0.00167; Fig 2E), whereas significantly lower settlement occurred in red wavelengths (*p*=0.01137; Fig 2E), compared to the other two wavelengths. No significant interaction between light intensity and light wavelength was detected.

### Multisensory integration and a sensory cue hierarchy determine settlement in actinulae

Multisensory integration is detected by examining statistical interactions between different sensory conditions (Stein et al. 2009). Here we examined interactions between sensory cues as they relate to settlement rate using a three-way ANOVA and contrasts based on our experimental approach (Fig 2A; Table S4). No interaction between light and chemical cues was identified across experiments (*p*=0.33; 70 experiments total; Fig 2F and Fig 2I). Similarly, no significant interaction (*p*=0.89) was observed between light and mechanical cues (Fig 2G and Fig 2J). Conversely, chemical and mechanical cues displayed a significant positive interaction (*p*=<2e-16; Fig 2K). Alone, chemosensory cues were a strong determinant of larval settlement. However, in the combined presence of the mechanical cue, settlement rates were significantly enhanced, indicating an inversion in the sign of the mechanical cue in the presence of the chemical cue (biofilm). Recall that when the chemical cue was absent, the mechanical cue had a significant negative influence on settlement rate (*p*=<2e-16; Fig 2K).

The three-way interaction between photo-, chemo-, and mechanosensory cues from the ANOVA is also significant (*p*=0.022) and indicates MSI by enhancement (Stein et al. 2009). The multisensory response has a significantly higher mean than the largest unisensory response (chemosensory) and the chemo-mechanical response in the absence of light (Fig 3; Fig 2H; Fig 2L). Among the three-way interaction, larval settlement was significantly lower in darkness when chemical and mechanical cues were present (*p*=0.03; Fig 2H and L). Additionally, the rate of settlement is significantly higher in green light in the presence of chemical and mechanical cues (*p*=0.026), and significantly lower in red light (*p*=0.003; Fig 2H and L). This interaction can be seen in Fig 2L, which shows that green light at both intensities induce higher settlement rates under chemical, and mechanical cues, and lower settlement rates occur in red light. Together, the Area under curve (AUC) data for the different combinations of cues depict a sensory cue hierarchy (Fig 3) where the highest rates of settlement occur in the presence of light, chemical, and mechanical cues.

**Figure 3.**
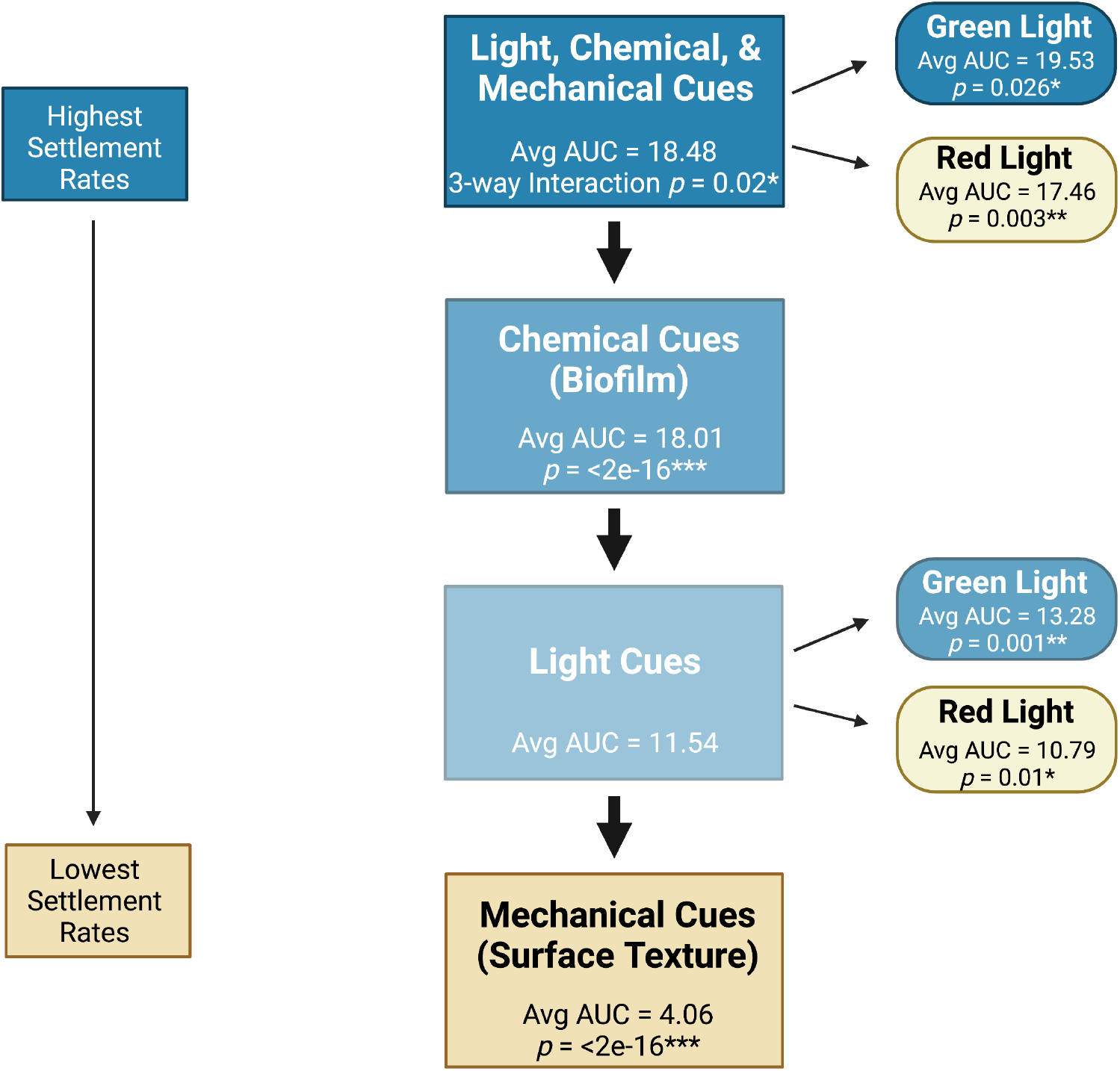
Sensory cue hierarchy of larval settlement in actinula larvae. The highest rates in settlement occur in the presence of all three cues, where significantly higher settlement rates occurred in the presence of green light, and significantly lower settlement rates occurred in the presence of red light. Following the combination of all three cues is the second tier composed of chemical cues which significantly increases larval settlement rates, but not to the same magnitude as having all three cues present. Light did not interact with chemical cues, leading to the lack of a light hierarchy in this second tier. The third tier is the light cues only treatment, where the highest rates of settlement occur in green light and the lowest rates occur in red light. Lastly, is the Mechanical cue tier which significantly decreased settlement rates. Additionally, the scale of color indicates the strength of a cue (darker colors indicate stronger influence).

### Sensory gene expression peaks in competent larvae and diminishes during and after metamorphosis

We performed a comparative transcriptome study on six developmental stages of *E. crocea*, from star embryo to metamorphosed juvenile polyp, to assess genetic correlates of the observed sensory response. Illumina Hi-seq sequencing resulted in an average of 38 million reads per replicate, with six replicates for each of the six developmental stages. We first used orthology analyses to place assembled *E. crocea* transcripts into orthogroups that included sequence data from a selection of metazoan genomes including human. We then used the Gene Set Enrichment Analysis (GSEA) database (Subramanian et al. 2005; Liberzon et al. 2011, 2015) and EdgeR to identify transcripts with significant differences in expression between any developmental stage that were present in three gene sets: Sensory perception of light stimulus (212 genes in gene set; Fig 4A), Sensory perception of chemical stimulus (501 genes in gene set; Fig 4B), and sensory perception of mechanical stimulus (164 genes in gene set; Fig 4C).

**Figure 4.**
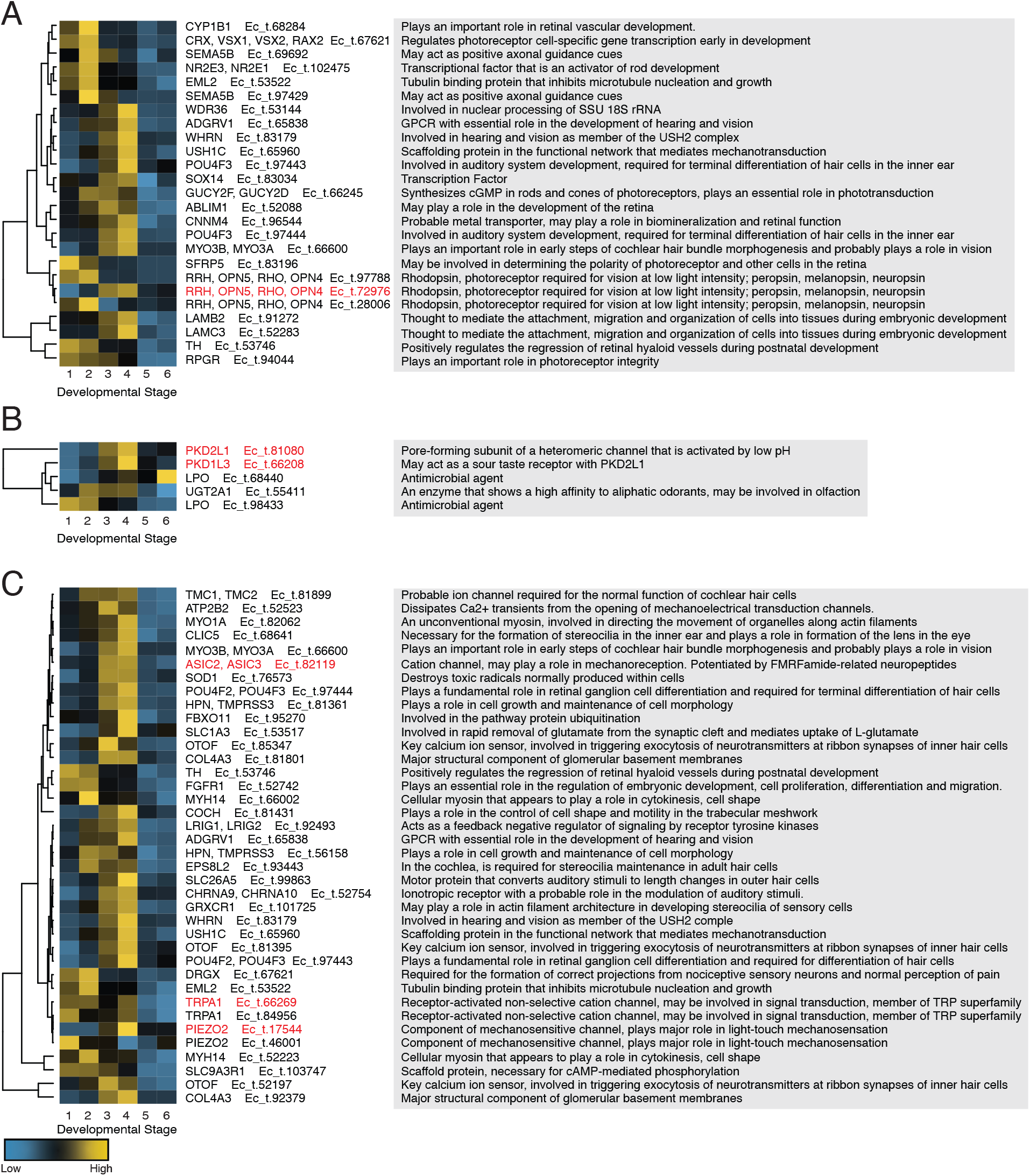
Gene expression over development in actinulae larvae for three sensory gene sets Heatmaps of significantly differentially expressed genes over development for the following three gene sets: (**A**) sensory perception of light stimulus; (**B**) sensory perception of chemical stimulus; and (**C**) sensory perception of mechanical stimulus. High expression is signified by yellow and low expression is light blue. The grey panels to the right contain abbreviated descriptions of gene functions from UniProt. The transcripts highlighted in red are the transcripts used to make RNA Fluorescent *in situ* hybridization probes. Gene trees of the orthogroups of probes can be seen in Figure S6. Developmental stages: 1 *star embryo*, 2 *preactinula*, 3 *young actinula*, 4 *actinula in nematocyte-printing stage*, 5 *settled (attached) actinula*, 6 *metamorphosed juvenile polyp*.

Gene expression for both the photo- and mechanosensory gene sets share similarities across larval development (Fig 4). First, genes involved in neurogenesis, cellular morphogenesis, and sensory cell type specification show maximum expression early and peak at stage 2 of development (preactinula). Examples of these for the sensory perception of light stimulus gene set include *E. crocea* homologs of SEMA5B (axonal guidance), homologs of CRX, VSX1, VSX2, and RAX2 (transcription factors, photoreceptor specification), NRL2E3 (transcription factor, rod specification), RRH (opsin, detection of light) and others (Fig 4A). A similar pattern is observed for the perception of mechanical stimuli gene set and include *E. crocea* homologs of MYH14 (cell shape and cytokinesis), EPS8L2 (stereocilia maintenance), DRGX (nociceptive sensory neuron development) and others (Fig 4C).

Next, a later pulse of gene expression associated with sensory physiology, cellular structure and morphogenesis, and another round of sensory cell type specification characterizes both gene sets. Examples of these later expressed transcripts for the perception of light stimuli gene set include homologs of GUCY2F and RRH (phototransduction), MYO3B (photoreceptor cell maintenance), ADGRV1 (development of hearing and vision), CNNM4 (retinal function), WHRN (periciliary membrane complex maintenance), and POU4F2 (retinal ganglion cell development). Genes that peak at stages 3 and 4 (actinula) from the perception of mechanical stimuli gene set include CLIC5 (stereocilia formation), MYO3B (cochlear hair bundle morphogenesis), POU4F2 (terminal differentiation of hair cells), SLC25A5 (motor protein, modulates auditory stimuli), ASIC2-3 (mechanoreception, potentiated by FMRFamide-related peptides), TRPA1 (mechanoreceptor transduction channel), and PIEZO2 (component of a mechanosensitive channel). Finally, transcript expression for both gene sets strongly diminishes at stages 5 and 6, which correspond to metamorphosis and the establishment of the primary polyp. This drop-off in expression is specific to the sensory gene sets used to interrogate our data and is not a general feature of our transcriptome data (Fig S3, Table S5). It is noteworthy that many of the transcripts that show differential expression in development for either the photo- or mechanosensory gene sets have shared functions in both.

Similar analyses for the sensory perception of chemical stimulus gene set indicate a markedly different trend as few developmentally differentially expressed transcripts were recovered for this set (Fig 4B). However, they do indicate strong stage 4 (actinula) differential expression of homologs of both PKD2L1 and PKD1L3, which dimerize and facilitate sour taste perception in mammals (Ishimaru et al. 2006; Fain 2020).

### RNA Fluorescent *in situ* hybridization reveal evidence for polymodal sensory cells

Our studies of larval nervous system development, settlement behavior, and developmental transcriptomics portray an inflection of sensory capacity at stage 4 when actinulae are competent to settle. In order to elucidate the cell types associated with enhanced sensory capacity, and potential cellular partitioning among the different sensory modalities, we conducted fluorescent *in situ* hybridization (FISH) using RNA probes against selected, highly differentiated sensory transcripts (Fig 4) in stage 4 *E. crocea* actinulae. Our targeted transcripts include opsin (Ec_t.72976, photoreception), Piezo and TRPA1 (Ec_t.17544 and Ec_t.66269 respectively, mechanotransduction), and ASIC, PKD2L1, and PKD2L3 (Ec_t.82119, Ec_t.81080 and Ec_t.66208 respectively, chemotransduction). We examined the cellular and spatial expression of these transcripts using the opsin probe as a reference present in all FISH experiments. Confocal optical sections (0.2 μm) indicate strong colocalization between opsin (Fig 5 A-I), PKD1L3 (Fig 5 A and C) PKD2L1 (Fig 5 D, F, G, I), and ASIC (Fig 5 A and B) in sensory neurons that are adjacent to cnidocytes in all experiments. Conversely, Piezo is strongly expressed in cnidocytes, with weaker expression of opsin also detected in cnidocytes (Fig 5 G-I). Finally, opsin and PKD2L1 show some co-expression with TRPA, but most TRPA expression is observed in cnidocytes to the exclusion of opsin and PKD2L1, which are expressed primarily in sensory neurons (Fig 5 D-F). Our results were confirmed by quantifying colocalization using Manders’ Overlap (Fig S4).

**Figure 5.**
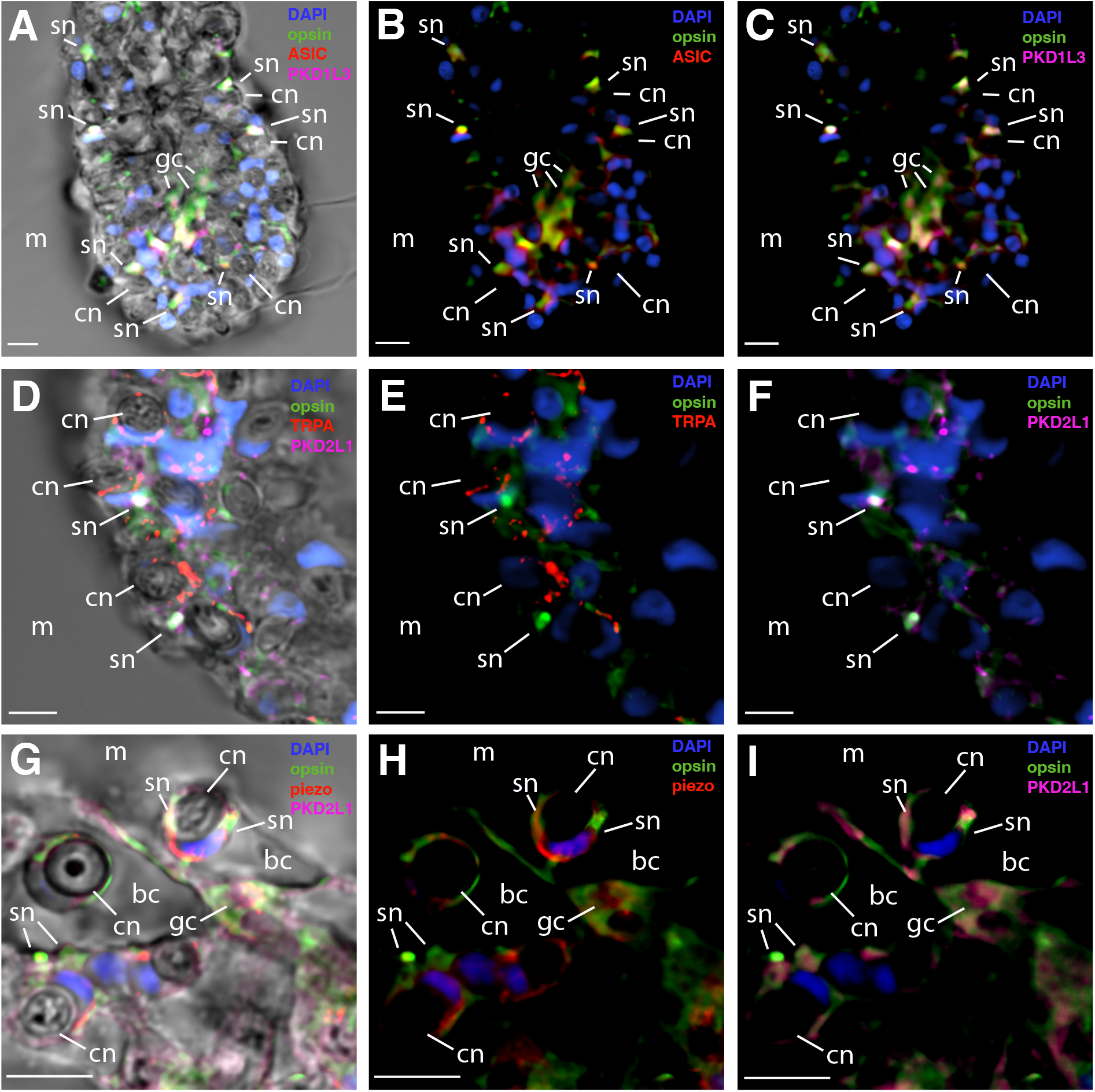
RNA fluorescent *in-situ* hybridizations (FISH) of sensory transcripts in actinulae tentacles (**A-C**) Same tissue where green is opsin, red is ASIC, cyan is PKD1L3, and blue is DAPI. (**A**) Merge of all confocal channels with the transmitted light (TD) channel. (**B**-**C**) The merge of confocal channels separated from transmitted light (TD). (**D-F**) Same tissue where green is opsin, red is TRPA, cyan is PKD2L1, and blue is DAPI. (**D**) Merge of confocal channels with the transmitted light (TD) channel. (**E**-**F**) The merge of confocal channels separated from the transmitted light (TD). (**G-I**) Same tissue where green is opsin, red is Piezo, cyan is PKD2L1, and blue is DAPI. (**G**) Merge of confocal channels with the transmitted light (TD) channel. (**H**-**I**) The merge of all confocal channels separated from the transmitted light (TD) at different sections of the sample. Cnidocytes (cn), sensory neurons (sn), ganglion cell (gc), battery cell (bc), medium (sea water) (m). All scale bars are 10um.

## Discussion

### Competence to settle is associated with cnidocyte and nervous system maturation

Our immunohistochemical analysis demonstrates that actinula (stage 4) larvae possess an elaborated nervous system with an abundance of cnidocytes located at the tips of the aboral tentacles and the basal protrusion. This is consistent with the findings of Yamashita et al. (2003), who described the cnidocyte composition of actinula larvae and showed that atrichous isorhizas are used to temporarily attach to surfaces in a behavior referred to as nematocyte-printing. In hydrozoans like *E. crocea*, cnidocyte discharge is modulated by adjacent sensory neurons, with which they form synapses (Hobmayer et al. 1990; Anderson et al. 2004; Westfall 2004; Anderson and Bouchard 2009; Plachetzki et al. 2012). The cnidocytes that facilitate nematocyte printing are located in the tentacles and a ring of neurons is present at the basal protrusion located on the aboral end of actinulae (Fig 1D). The aboral region is also the site for sensory integration in other hydrozoan larvae (Matsushima et al. 2010; Tran and Hadfield 2013). We propose that the sensory interplay between cnidocytes and their adjacent sensory neurons facilitate sensory integration during settlement.

### A sensory cue hierarchy and MSI during the larval settlement decision

Our larval settlement studies identified a sensory cue hierarchy, where the highest rates of settlement occurred in the presence of chemical, mechanical, and light cues, with green light being the most permissive to settlement (Fig 3). This is not surprising as marine invertebrate larvae in general are known to integrate information from different modalities (Pawlik 1992; Kawaii et al. 1997; Hadfield and Paul 2001; Holst and Jarms 2007; Hadfield 2011; Mason and Cohen 2012; Strader et al. 2015; Whalan et al. 2015; Say and Degnan 2020). This type of sensory integration, and the likelihood that some cues are more important than others, was the basis for the proposal that a hierarchy of sensory cues dictates larval settlement in a species-specific manner (Kingsford et al. 2002; Hodin et al. 2018). However, the existence of such a hierarchy has not been experimentally validated until now. At the top of the sensory cue hierarchy for *E. crocea* larval settlement are chemical cues produced by microbial biofilms. This is consistent with larval settlement studies in other metazoan species (e.g., annelids: (Hadfield and Paul 2001; Harder et al. 2002; Lau et al. 2002; Huang and Hadfield 2003; Huang et al. 2007; Hadfield 2011; Freckelton et al. 2017); barnacles and bryozoans: (Qian et al. 2003; Faimali et al. 2004; Dobretsov and Qian 2006; Zardus et al. 2008; Dobretsov and Rittschof 2020)) that identified biofilm-derived cues for larval settlement. It is likely that biofilm-derived chemical cues serve as a general indicator of habitat quality across sessile metazoan species with dispersive larval stages (Wieczorek and Todd 1998; Lau et al. 2005; Hadfield 2011).

We also found a significant effect of light wavelength on larval settlement in *E. crocea*. Previous work in hydrozoan species noted similar findings where animals are more sensitive to light in the blue to blue–green spectrum compared to red light (Taddei-Ferretti et al. 2004; Guertin and Kass-Simon 2015; Strader et al. 2015). In the case of *E. crocea* larval settlement, green light, which penetrates only to shallow depths in the water column, may serve as a depth meter ensuring that colonies of *E. crocea* settle at shallow depths where prey items like plankton are abundant. We note that our finding of the preference of *E. crocea* larvae to settle in areas illuminated in green light does not in itself indicate the capacity for discrimination between wavelengths of light in *E. crocea* larvae. It is more likely that actinulae larvae are effectively “color blind” but show a higher sensitivity to green light due to a limited photoreceptor palate with sensitivity in that range. Indeed, we uncovered only two closely related opsin transcripts from *E. crocea* larval stages.

Many of the same sensory cues that we investigated here have been investigated previously in other species. However, in most cases, studies have focused on one or two cues of interest (Svane and Dolmer 1995; Nellis and Bourget 1996; Wieczorek and Todd 1998; Zardus et al. 2008; Mason and Cohen 2012; Strader et al. 2015; Hodin et al. 2018; Say and Degnan 2020). Our factorial experimental design allowed us to identify interactions between sensory modalities and to test the possibility of MSI. Surprisingly, we found a significant interaction between mechanical (surface texture) and chemical (biofilm) cues whereas, in the presence of chemical cues, surface texture enhances settlement rates, but in the absence of chemical cues, surface texture is inhibitory. This significant and sign-reversing sensory interaction constitutes MSI by enhancement (Stein et al. 2009) and is likely mediated by signaling between sensory neurons and their adjacent cnidocytes. MSI by enhancement is stronger when light is present in addition to chemical and mechanical cues, suggesting that both photosensitivity and chemosensitivity have positive impacts on settlement when mechanical cues are present.

### The cellular basis for MSI in actinulae

Temporal patterns of gene expression also illuminate the sensory determinants of the settlement decision in *E. crocea*. Our analytical approach, a type of GSEA, filtered transcripts based on their phylogenetic membership into clades containing genes annotated as having sensory function and then interrogated such transcripts based on their differential expression between stages. While each of the three gene sets used for interrogation contained greater than 200 genes, only a subset of those genes have orthologs in the *E. crocea* transcriptome and are differentially expressed between stages (Fig 4). The mechanosensory gene set includes the largest number of active genes while the chemosensory gene set contains the least number of genes. This disparity is partly due to the relative degree of conservation in gene function between cnidarian genomes and those of model organisms from which annotations are based, and partly due to genes that are differentially expressed in *E. crocea* larval development. In addition, the mechanosensory and photosensory gene sets share a number of genes in common due to pleiotropic expression. It is somewhat surprising that so few chemosensory genes were recovered by our screen given that chemosensitivity is the predominant sensory cue in larval settlement. However, we did identify orthologs of both PKD1L3 and PKD2L1, which have been implicated as key components of the sour taste (pH) transduction pathway in mammals (Ishimaru et al. 2006; Fain 2020) and have previously been implicated in cnidarian chemosensitivity (McLaughlin 2017). Furthermore, biofilms vary in acidity (Dexter and Chandrasekaran 2000), which could allow for the assessment of different settlement sites based on their chemosensory properties.

These analyses identify two pulses of sensory gene expression: one occurring early (stage 2, preactinula) and consisting largely of regulatory and structural factors and another occurring later (stage 4, actinula) and consisting of structural and physiological factors. These data add further support for the actinula (stage 4) as the maximally sensory-equipped larval stage and highlight candidate genes for expression analyses.

We examined the expression of several candidate genes in stage 4 actinula larvae using FISH (Fig 5). We used the same opsin transcript (Ec_t.72976) in each experiment to facilitate comparisons between genes. Our results demonstrate strong co-expression of opsin, PKD2L1, PKD1L3 and ASIC in sensory neurons. In contrast, TRPA and Piezo transcripts localized to cnidocytes. These results demonstrate cellular partitioning between the modes of sensation, where opsin (photosensitivity) and PKD2L1, PKD1L3, and ASIC (chemosensitivity) are expressed in polymodal sensory neurons, and TRPA and Piezo (mechanosensitivity) are expressed in cnidocytes.

Together, these data suggest that MSI in *E. crocea* larval settlement is facilitated by a simple communication circuit between polymodal photo-chemo sensory neurons and mechanoreceptive cnidocytes located on the tentacles and aboral basal protrusion. We propose that in the absence of chemical and photosensory cues, mechanosensitive cnidocytes are inhibited from discharging and little cnidocyte printing behavior takes place. However, when light and chemical cues are present, this inhibition is relieved and mechanosensitive cnidocytes are free to fire. The combination of permissive light, chemical, and mechanical cues lead to the highest rate of settlement.

MSI is best known in animals with complex brains where specialized brain centers have evolved to facilitate information processing and exchange. Here we show that MSI is possible in animals that lack centralized nervous systems and may be facilitated by communication between as few as two cell types, sensory neurons and cnidocytes. Moreover, given the importance of larval settlement dynamics in shaping benthic ecosystems, it is likely that MSI as observed in *E. crocea* larvae, may be an important determinant of benthic community composition and function.

## Conclusion

Understanding how cnidarians integrate sensory information from the environment is critical to understanding the ecological processes that dictate benthic community composition. At the same time, uncovering the genetic determinants of cnidarian sensory behavior can illuminate the deep evolutionary histories of the animal senses and provide clues on their early functions. We show that brainless actinula larvae use multisensory integration (MSI) during the larval settlement decision that incorporates information processing from the light, chemical and mechanical senses. MSI is usually portrayed as a process involving information flow between higher-level brain centers (Stein 1998; Otto et al. 2013; Stein et al. 2014; Ghosh et al. 2017; Currier and Nagel 2020), however, our results indicate that MSI may be facilitated by interactions between cells, and may have been a prominent feature of the organismal biology of metazoans prior to the evolution of complex brains.

## Acknowledgments

Research reported in this publication was supported through the University of New Hampshire’s Center for Integrated Biomedical and Bioengineering Research (CIBBR) through a grant from the USDA.

## Author Contributions

Conceptualization, D.P.; Methodology, D.P. and S.B.; Investigation, S.B.; Writing – Original Draft, S.B.; Review & Editing, D.P. and S.B.; Funding Acquisition, D.P.; Resources, D.P.; Supervision, D.P.

## Methods

### Field collection of *E. crocea* Colonies

Larva were obtained by collecting adult *E. crocea* colonies at the UNH Coastal Marine Lab (CML) pier in New Castle, NH in June and July 2020. Colonies were transported to UNH in unfiltered sea water immediately after collection. The colonies were cultured and maintained according to Mackie (1966) where colonies were placed in large glass bowls for predator and dirt removal. Once cleaned, colonies were moved to new bowls filled with unfiltered sea water and were placed in an incubator with an aeration stone connected to a bubbler at a 12/12 L:D cycle at 18°C. Colonies were undisturbed overnight to allow spawning and actinula release. Actinula were identified and collected the following day for settlement experiments or for staining.

### Immunohistochemistry and Confocal Microcopy

Immunohistochemistry was performed on four developmental stages: embryos, pre-actinulae, actinula larvae, and juvenile polyps, which were obtained as described above. Samples were fixed overnight at 4°C in 4% paraformaldehyde (Sigma) in PBST. We followed the protocol from Plachetzki et al. (2012) with minor alterations for staining. Samples were washed five times with 5 minute incubations in PBST (3.2 mM Na2HPO4, 0.5 mM KH2PO4, 1.3 mM KCl, 135 mM NaCl, 0.1% Tween 20, pH 7.4) and blocked for 2 h in PBST + 20% normal goat serum (NGS; Sigma) at room temperature. Samples were then incubated with primary antibody, anti-acetylated *α* tubulin (1:500; Sigma), in blocking solution over night at 4°C. Following the primary antibody, samples were washed five times with 5 min incubations in PBST and blocked as before. Samples were then incubated with the secondary antibody, Cy2-conjugated anti mouse IgG (1:1000; Jackson), and blocking reagent overnight at 4°C. Samples were then washed five times with 5 min incubations in PBST. Afterwards, a solution containing Alexa Fluor 488-labeled phalloidin stock (1:40; Invitrogen) in PBST was added for 1 h. Samples were then washed five times with 10 min incubations and were then mounted in ProLong Antifade Mountant with DAPI (ThermoFisher). Samples were imaged on the UNH Nikon A1R HD confocal Microscope.

### Larval Settlement Study Experimental Design and Materials

The larval settlement study was designed as a 7×2×2 split-plot Randomized Complete Block Design (RCBD) with 10 blocks, Fig 1A. The blocking variable was the experimental day that replicates were performed on where larvae in one block were from colonies collected at the same time and location the day prior to the experiment. The purpose of the blocking variable was to account for environmental variation between colonies and days across the study, and each block contained all 28 treatments. An experimental unit was a single petri dish that contained 10 actinula larvae, this allowed us to use the area under the curve (AUC) as our response variable. The three treatment factors were the sensory cues of interest which included seven levels of light conditions, two levels of chemosensitive treatments, and two levels of mechanosensitive treatments. Replicates were performed in four hand-made light boxes where each light box had two chambers (described below). The first randomization of the split-plot design, which was applied at the chamber level, were the seven light conditions described in Table 1. The second randomization was comprised of the chemical and mechanical cues and were applied at the petri-dish level in a factorial presence/absence structure and can be seen in Table 2.

**Table 1.** The seven levels of the light condition factor with their corresponding measurements

| Light Conditions in Study | Wavelength | Intensity |
| --- | --- | --- |
| Red Light, High Intensity | 630nm | 10 $\mu\text{mol m}^{-2} \text{s}^{-1}$ |
| Red Light, Low Intensity | 630nm | 5 $\mu\text{mol m}^{-2} \text{s}^{-1}$ |
| Green Light, High Intensity | 520 nm | 10 $\mu\text{mol m}^{-2} \text{s}^{-1}$ |
| Green Light, Low Intensity | 520 nm | 5 $\mu\text{mol m}^{-2} \text{s}^{-1}$ |
| Blue Light, High Intensity | 460nm | 10 $\mu\text{mol m}^{-2} \text{s}^{-1}$ |
| Blue Light, Low Intensity | 460nm | 5 $\mu\text{mol m}^{-2} \text{s}^{-1}$ |
| Darkness | 0nm | 0 $\mu\text{mol m}^{-2} \text{s}^{-1}$ |

**Table 2.** Factorial Structure of Chemical and Mechanical Cue Dish Treatments

|  | Chemical Cue Absent | Chemical Cue Present |
| --- | --- | --- |
| Mechanical Cue Absent | Dish Treatment A | Dish Treatment B |
| Mechanical Cue Present | Dish Treatment C | Dish Treatment D |

Prior to settlement experiments, four five-sided light boxes were created with black opaque acrylic (Acrylite; Extruded 9M001) with a thickness of 0.177 inch (in) and 16in x 12in x 12in dimensions with two centered, circular holes with diameters of 85mm cut out of the top. An additional piece of acrylic with the same thickness with 12in x 12in dimensions was placed in the center of the box to create two separate chambers. Super Bright LEDs heatless lights (part #: WRLFA-RGB6W-60) were placed in the cutout holes which allowed control of four different light intensity settings for the wavelengths red (630 nm), green (520 nm), and blue (460 nm). Light intensity was measured using an LI-1000 DataLogger. For replicability the boxes were placed in the same position for each replicate. The positions of petri-dishes were also marked at high and low intensities for each wavelength in each chamber for replicability, Fig S1B.

To obtain chemical cues, we allowed petri-dishes to generate biofilms by placing dishes in mesh dive bags and attached them to the UNH CML pier for 1 week at a depth of 1.5 m (Lee et al. 2008, 2014). Colonized dishes were then transported to the laboratory in seawater collected at the site and were used in settlement studies and for biofilm sequencing (Corcoll et al. 2017). To obtain mechanical stimuli, petri-dishes were scratched by 36-Grit Ceramic Alumina Sandpaper (Lowes; Model #: 9150-052) in a circular motion on the outer part of the dish, then with three scratches, non-overlapping, going from one side to the other in the center of the dish to ensure full coverage. Dishes were then rinsed twice, once with DI water to remove any plastic shavings, and then a second time with sea water immediately prior of use in a replicate to ensure removal of plastic shavings.

### Larval Settlement Study

Larva were obtained by collecting adult *E. crocea* colonies at the UNH CML pier in New Castle, NH in June and July 2020. Colonies were cultured and maintained according to Mackie (1966) as described above. Colonies of *E. crocea* were undisturbed overnight (12-15 hours) to allow spawning and actinula release. The following day, larvae at the actinula stage were collected and placed in 100mm × 15mm plastic petri dishes (Thermo Scientific) where each petri dish contained 10 larvae and the dish treatments corresponded to the predetermined randomized dish conditions for the 28 experimental treatments. Larvae were identified under a microscope following Yamashita et al. (2003) where we sought out young actinula with stiff tentacles, small and circular bodies, and short aboral poles.

One block (replicate) of all 28 treatment conditions began once larvae were collected at the actinula stage where start times were staggered by chamber which had four petri dishes with a total of 40 larvae. Dishes from the four chemical and mechanical treatment conditions were placed at the appropriate positions and wavelength and light intensity were adjusted according to the predetermined randomized treatment conditions per chamber. Metamorphic stages were recorded at the following time points: 0 hours (hrs), 2hrs, 4hrs, 6hrs, 8hrs, 12hrs, and 24hrs. At each time point, dishes were taken out of the light box and placed under a microscope where each individual larva was observed and quantified based on their metamorphic stage (detailed below). Immediately following quantification, the replicate dish was placed back into the light box until the next time point for quantification, this procedure was performed for each of the 28 dishes at each time point.

Larval quantification was assessed on a presence/absence metamorphosis scale with three levels. A 0 signified larvae were still in the actinula phase and had not metamorphosed or settled, a 1 indicated that larvae had settled (attached to the substrate but had not completed metamorphosis), and a 2 signified that larvae had completed metamorphosis into a juvenile polyp. At each time point, we counted and recorded the total number of larvae for each condition under the scale. From this information, we then calculated the area under the curve using the settlement percentages at each time point, which combined the number of settled and metamorphosed larvae (# of Settled + # of Metamorphosed/total # of larvae in dish). The following equation was used to calculate the area under the curve (AUC) which was used as the response variable in statistical analyses (Mohapatra et al, 2014; Shaner, 1977):

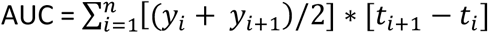

Where *y*_*i*_ is the proportion of settled and metamorphosed larvae at the *i*^th^ time point; *t*_*i*_ is the timepoint in hours where larvae were observed, and *n* is the total number of observations per petri-dish in a replicate. Normally distributed data were compared statistically by a split-plot RCBD three-way analysis of variance (ANOVA) and with orthogonal contrasts in the R environment.

### Library preparation, sequencing and read processing

Colonies of *E. crocea* were collected at the UNH CML pier in May and June 2019. Colonies were maintained and larva were obtained as previously mentioned above. We collected six replicates of the six developmental stages: embryos, pre-actinula, actinula (younger), settling actinula (older), settled actinula, and metamorphosed juvenile polyps (Yamashita 2003), where each stage had a total of 125 larva collected. Samples were stored in RNA*later* (Thermo Fisher Scientific) in −20°C until total RNA was extracted using the PureLink RNA Mini Kit (Thermo Fisher Scientific; Cat no. 12183018A) according to the manufacturer’s instructions and quantified using NanoDrop (Thermo Scientific). Libraries were created using 1000ng of total RNA with the NEBNext Ultra 2 Directional RNA Library Kit following the Poly(A) mRNA Magnetic Isolation module (NEB #E7490). Libraries were sent to Novogene for Illumina Hi-Seq. After sequencing, reads were processed using custom python scripts which executed FastQC and selected the highest quality replicate for each of the 6 stages. It then concatenated the highest quality replicate for each of the 6 stages to generate representative R1 and R2 read files to be used in *de novo* transcriptome assembly. The Oyster River Protocol (ORP) was used as the assembler which performs read trimming, read normalization, read error correction, and generates an assembly using a multi-kmer and multi-assembler approach and then merges those assemblies into one final high-quality assembly (MacManes 2018). The ORP also produced quality metrics from TransRate and BUSCO.

### Gene expression analysis

To prepare for gene expression analyses performed later in the workflow, two sub-steps were completed. First, the program Salmon was executed to quantify transcripts in the linux environment (Patro et al., 2017). Next, EdgeR was implemented in the R environment to identify significant differential gene expression between pair-wise comparisons of the six developmental stages (Chen et al. 2020). Count data was reduced by requiring transcripts to have 10 counts in a library to be considered expressed and were filtered by counts-per-million (CPM)(Chen et al. 2020). The calcNormFactors function was used to normalize library sizes which used a trimmed mean of M-values (TMM) method. To estimate dispersion, a quantile-adjusted conditional maximum likelihood (qCML) method was employed. Differentially expressed genes (DEGs) were identified at a p-value of 0.05 and results from each comparison was exported to the terminal and included the transcript header, logFC, logCPM, PValue, and FDR which was used in a later step.

TransDecoder was then implemented to translate the assembly from nucleotide space into protein space. To reduce the amount of duplicate protein models, we ran cd-hit on the actinula protein FASTA. The reduced actinula FASTA was then used in OrthoFinder along with the following taxa from publicly available data on NCBI: *Homo sapiens, Drosophila, Hydra, Hydractinia* (adult and larval forms), and *Nematostella*.

While OrthoFinder was running, we obtained sensory gene sets from the Gene Set Enrichment Analysis (GSEA) created by the Broad Institute and UC San Diego (Subramanian et al. 2005; Liberzon et al. 2011, 2015). We selected the following three gene sets: GO_Sensory_Percep_Of_Light_Stimulus, GO_Sensory_Perception_of_Chemical_Stimulus, and GO_Sensory_Perception_of_Mechanical_Stimulus. We then imported the Entrez IDs, gene symbols, and descriptions of the members in the gene sets as a CSV file into the terminal. Using custom made python scripts we reformatted the gene set information and searched the Entrez IDs using the Bio.Entrez module in python to obtain the protein sequences for each gene across the three gene sets.

Our human protein sequences used in OrthoFinder contained altered headers, therefore we performed three BLASTp searches on our human protein sequences using the genes from the sensory gene sets as queries. The search allowed one target sequence result at an e-value of 0.00001. The output from the three searches included the official NCBI headers and the corresponding altered header used in our human protein models. This was used to annotate the altered human protein models used in OrthoFinder with the various sensory genes in the following step. Additionally, we generated another output file that was composed of human sequence IDs from the NCBI database and their corresponding gene symbols.

To identify actinula transcripts that shared orthogroups with human transcripts of sensory genes, we imported the following results into the R environment to be used in a custom R script: OrthoFinder (orthogroup.tsv file and counts file), the BLAST results with the two human headers (NCBI and the altered headers used in our human protein FASTA), and the file with the NCBI headers and its corresponding gene symbol. This R script then identified and re-labeled the orthogroups that contained the genes from the sensory gene sets with the gene symbols and reported the actinula transcripts that were contained in those orthogroups with the gene symbol labels. The lists of actinula headers with their corresponding gene symbols were exported from R into the terminal and were used with a custom python script and the edgeR output to identify significantly differentially expressed genes within the gene sets. The significant DEGs were then visualized as heatmaps for each gene set in the R environment.

### RNA Fluorescent *in situ* Hybridization Study

Sequences were identified to make RNA probes for *in situ* hybridizations using the transcriptomic expression data from the developmental transcriptome study. We identified the highest expressed transcripts of key sensory genes from the three sensory gene sets. We then designed RNA-probes for our target sequences using the Stellaris RNA FISH platform (Biosearch Technologies) with the custom probe design service following their recommendations for probe design.

Actinula larvae were obtained as previously described above. Samples were stained according to the Stellaris FISH protocol with alterations to the protocol. Samples were fixed overnight at 4°C in 4% paraformaldehyde (Sigma) in PBS. The following day samples were washed five times with 5 minute incubations in PBST. Samples were then incubated in Prot K (1ug/ul) for 10 minutes at room temperature. Immediately following the Prot K incubation, the solution was removed, and samples were incubated in glycine (4ug/ul) for 10 minutes at room temperature to stop the Prot K reaction. Samples were then washed twice with 5 minute incubations in PBST. We then refixed samples in 4% PF for 30 minutes at room temperature. Following refixation, samples were washed five times with 5 minute incubations in PBST. We then transferred samples to 1.5ml microcentrifuge tubes and incubated samples in Wash Buffer A (Catalog# SMF-WA1-60) for 5 minutes at room temperature. We then ran a prehybridization step using the stellaris hybridization buffer (Catalog# SMF-HB1-10) at 37C for 1 hour in a shake ‘N’ bake Hybridization Oven. Following the prehybridization step, we added the hybridization buffer with probes (10ul of each probe for a total of 30ul of probe) to our samples and incubated them at 37C overnight in the dark. The following day, the hybridization buffer with probe solution was removed and samples were incubated in Wash Buffer A at 37C for 30 minutes in the dark. Samples were then incubated in Wash buffer B (Catalog# SMF-WB1-20) for 5 minutes at room temperature in the dark. We resuspended samples in PBST and mounted them in ProLong Antifade Mountant with DAPI (ThermoFisher). Samples were imaged on the UNH Nikon A1R HD confocal Microscope.

## Supplemental Information titles and legends

**Figure S1**. Experimental setup for actinula settlement study

**(A)** The lightboxes used in the experiment. Each lightbox had two separate chambers where each chamber had its own light source which could be changed to specific wavelengths of light and intensity as demonstrated in (**B**).

**Figure S2**. Block interactions of actinula settlement study

(**A**) The interaction between chemical cues and the blocking variable (experimental day). There is a significant interaction (*p*=0.002) occurring, however this figure depicts that the magnitude of the response to chemical cues is maintained over time (the blocking variable). (**B**) The interaction between mechanical cues and the blocking variable (experimental day). There is a significant interaction (*p*=0.005) occurring, however the magnitude of the two responses is the same, the lines do not intersect, which indicates that the blocks are not influencing the treatments.

**Figure S3**. Word Clouds of GO terms from significantly upregulated genes in actinula stages 5 and 6

(**A**) The GO terms for the significantly differentially expressed genes upregulated in stage 5, settled larvae. Created using ReviGO. (**B**) The GO terms for the significantly differentially expressed genes upregulated in stage 6, metamorphosed juvenile polyps. Created using ReviGO. Also see Table S5 for a list of significant GO terms.

**Figure S4**. Quantification of Fluorescent *in-situ* Hybridization colocalization in actinula larvae using Manders’ Overlap

To quantify colocalization in the fluorescent *in-situ* Hybridization study, we performed a Manders’ overlap analysis which identifies the percentage that pixels overlap. (**A**) Depicts the region of interest (ROI) where the Manders’ overlap was performed for the staining of Opsin (EGFP), Piezo (RFP), and PKD2L1 (CY5). The first row in the table details the percentage of overlap for this experiment for each channel comparison. (**B**) Depicts the region of interest (ROI) for the staining of opsin (EGFP), TRPA (RFP), and PKD2L1 (CY5). The second row in the table details the percentage of overlap for this experiment. (**C**) Depicts the region of interest for the staining of opsin (EGFP), ASIC (RFP), and PKD1L3(CY5). Across all experiments, DAPI stained cell nuclei blue. All scale bars are 10um.

**Figure S5**. Control for RNA Fluorescent in-situ Hybridizations

To confirm that the staining from Figure 5 was actual signal, we performed a control experiment where samples underwent the same protocol apart from adding RNA probes. A-J Two images of different control samples. (A&F) Merged images of all the channels. (B&G) The RFP channel. (C&H) The CY5 channel. (D&I) The GFP channel. (E&J) DAPI staining cell nuclei in blue.

**Figure S6**. Orthogroup gene trees of sequences used as RNA FISH Probes

The gene trees from Orthofinder for each of the 6 RNA FISH probes. Red indicates the transcript used as a probe. Green indicates an actinula transcript that was significantly differentially expressed (DE). Blue and the other colors represent the annotated human sequences from the gene set.

**Table S1**. Three-way ANOVA from actinula larva settlement study (7×2×2 split-plot RCBD)

The main plot (first randomized variable) is the light condition and the sub plot (second randomized variable) is the dish treatments which are set up in a factorial structure of presence/absence of chemical (biofilm) and mechanical (surface texture) cues.

**Table S2**. Block interaction investigation from actinula larva settlement study (7×2×2 split-plot RCBD)

We probed the block interaction because the blocking variable in Table S1 was significant. This ANOVA examines if the blocking variable interacts with the treatment variables.

**Table S3**. Photosensory contrasts in light-only treatment from the actinula larva settlement study Three separate contrasts were run, one for each wavelength of light, and are grouped together in this single table. We asked four questions: Is there a difference in larval settlement when larvae are (1) exposed to light compared to darkness; (2) in high-intensity light compared to low-intensity light? (3) Is there a difference in larval settlement when larvae are in green light compared to Blue & Red (Blue vs Green & Red, and Red vs Blue & Green); and (4) Is there a difference in settlement when larvae are exposed to green light at a high intensity vs low intensity (in blue light at high vs low; and red light at high vs low).

**Table S4**. Photosensory Contrasts for all interactions from the actinula larva settlement study Three separate contrasts were run, one for each wavelength, and are grouped together. Here we asked the same photosensory questions but are probing all two-way and the three-way interactions.

**Table S5**. Actinula stages 5 and 6 significantly upregulated GO terms used in ReviGO analysis (Supek et al. 2011).

We identified all genes that were significantly upregulated across these two stages and ran Interproscan to identify GO terms. These are the resulting GO terms that we input to ReviGO to generate word clouds of potential functions.

