## Supplementary Information for "Multisensory integration (MSI) by polymodal sensory neurons dictates larval settlement in a brainless cnidarian larva"

### I. Supplemental Figures

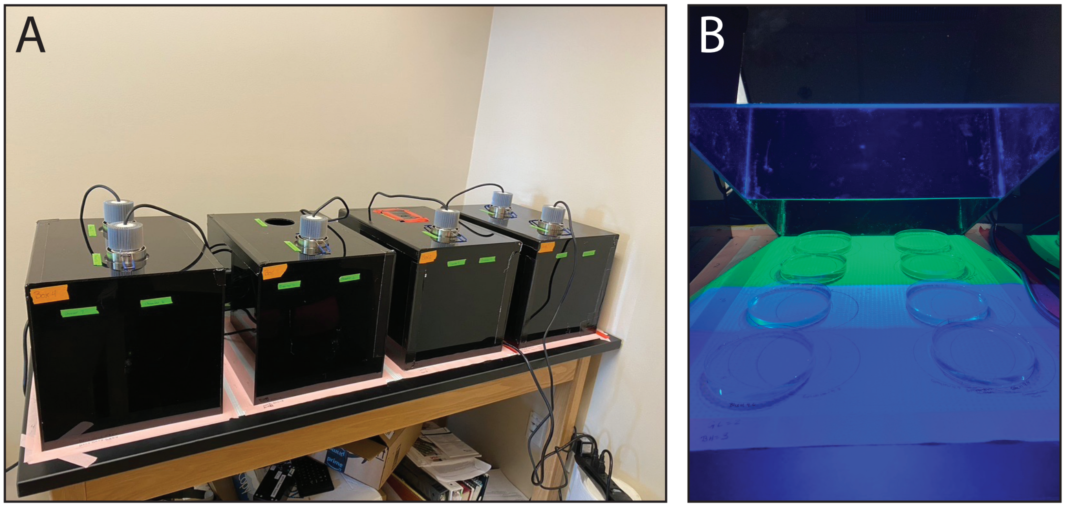

**Figure S1.** Experimental setup for actinula settlement study

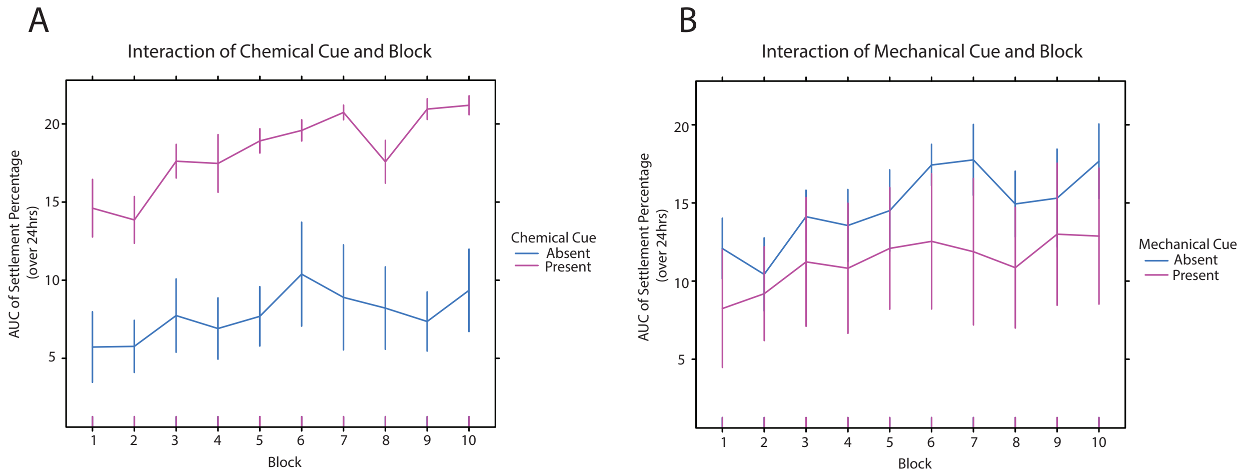

**Figure S2.** Block interactions of actinula settlement study

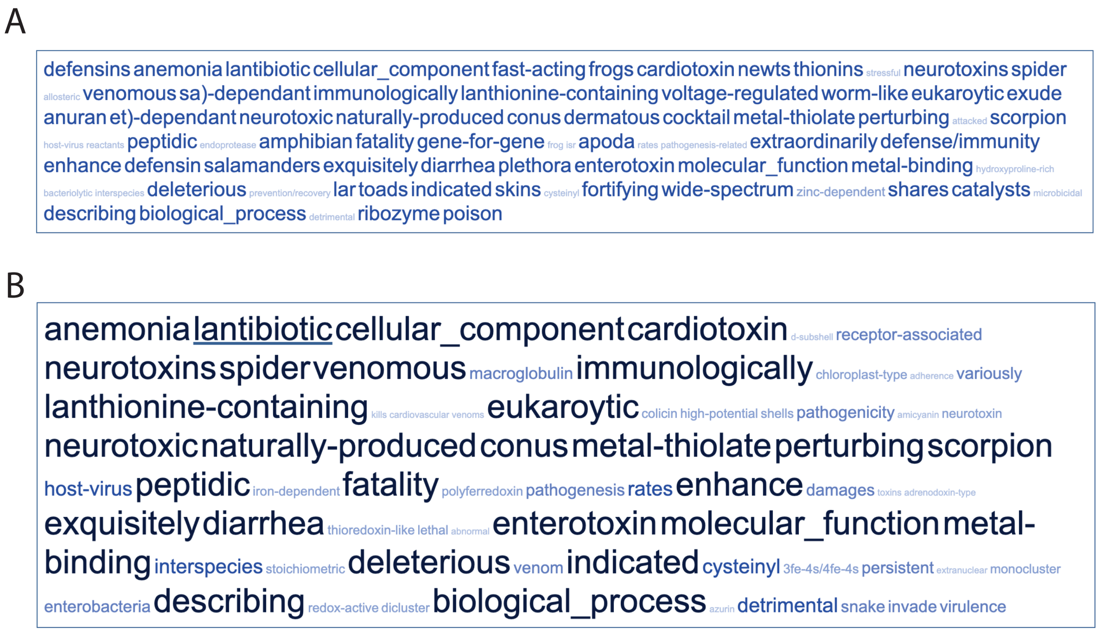

**Figure S3.** Word Clouds of GO terms from significantly upregulated genes in actinula stages 5 and 6

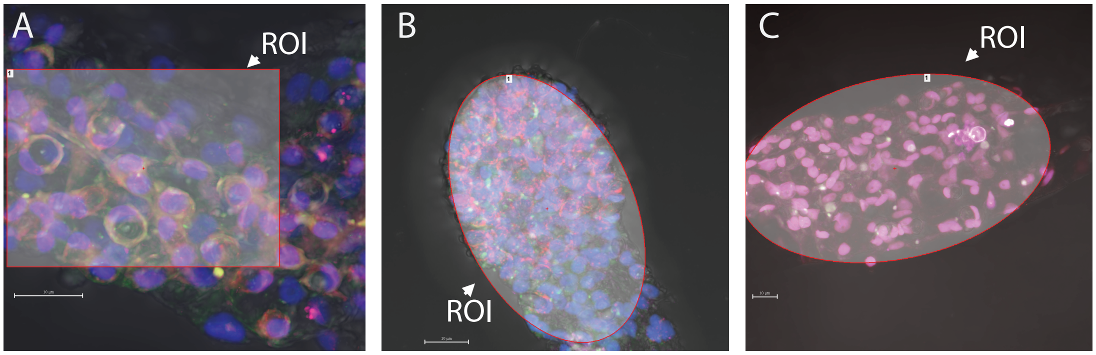

| COLOCALIZATION MANDERS' OVERLAP (GFP; RFP; CY5) | EGFP VS CY5 | EGFP VS RFP | CY5 VS RFP | EGFP VS DAPI | RFP VS DAPI | CY5 VS DAPI |
|---|---|---|---|---|---|---|
| (A) OPSIN; PIEZO; PKD2L1 | 0.99 | 0.97 | 0.97 | 0.70 | 0.80 | 0.70 |
| (B) OPSIN; ASIC; PKD1L3 | 0.98 | 0.89 | 0.90 | 0.77 | 0.85 | 0.79 |
| (C) OPSIN; TRPA; PKD2L1 | 0.99 | 0.86 | 0.86 | 0.50 | 0.85 | 0.50 |

**Figure S4.** Quantification of Fluorescent *in-situ* Hybridization colocalization in actinula larvae using Manders' Overlap

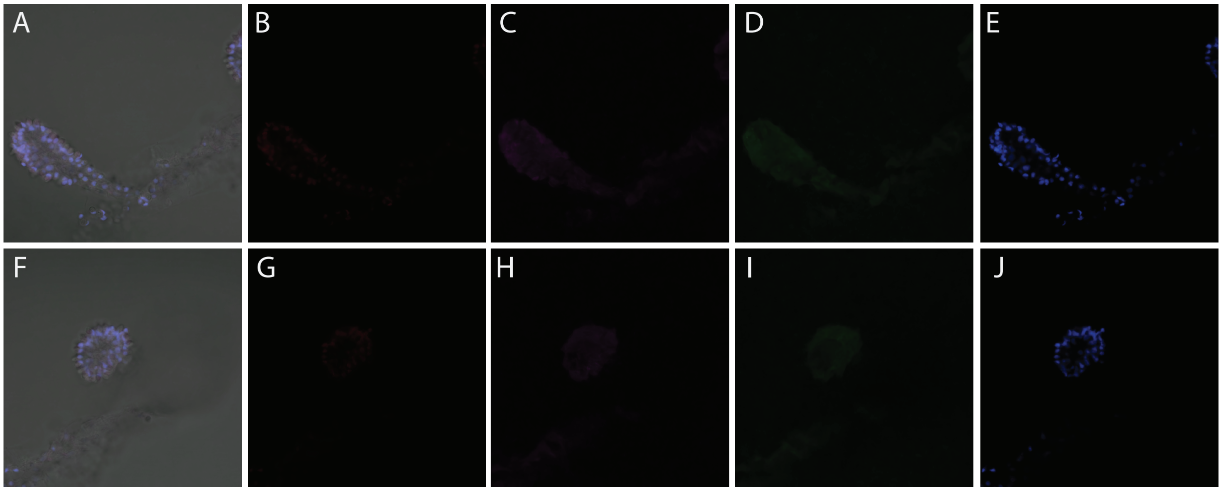

**Figure S5.** Control for RNA Fluorescent in-situ Hybridizations

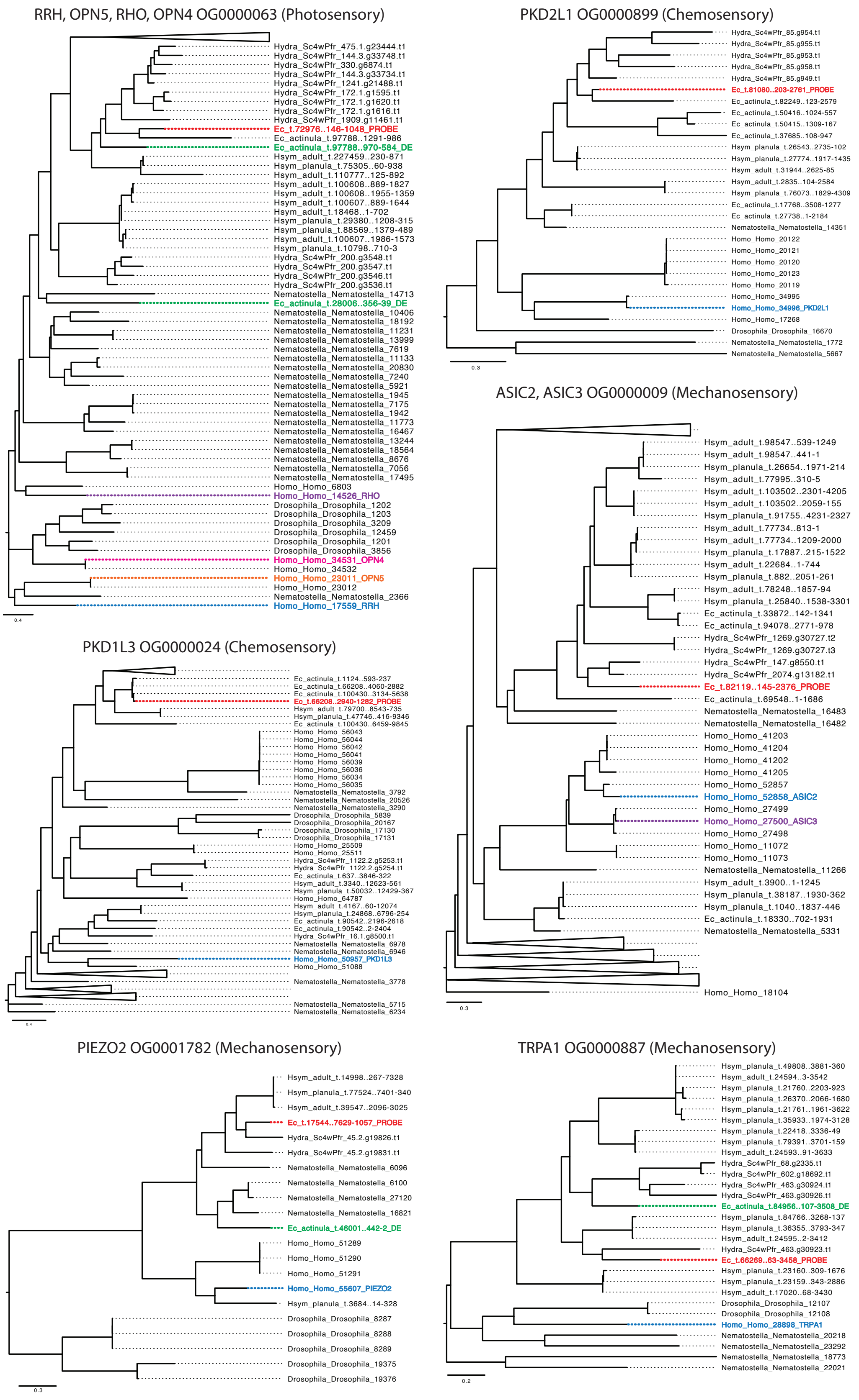

**Figure S6**. Orthogroup gene trees of sequences used as RNA FISH Probes.

### II. Supplemental Tables

| | DF | Sum Sq | Mean Sq | F-value | p-value |
|---|---|---|---|---|---|
| Light Condition | 6 | 83.3 | 13.88 | 2.293 | 0.0482 |
| Block | 9 | 918.6 | 102.07 | 16.858 | 8.66e-13 |
| Residuals (Light:Block) | 54 | 326.9 | 6.05 | | |
| Chemical cue | 1 | 7642 | 7642 | 1205.762 | <2e-16 |
| Mechanical Cue | 1 | 860 | 860 | 135.738 | <2e-16 |
| Light:Chemical | 6 | 43 | 7 | 1.39 | 0.3413 |
| Light:Mechanical | 6 | 14 | 2 | 0.373 | 0.8955 |
| Chemical:Mechanical | 1 | 1105 | 1105 | 174.323 | <2e-16 |
| Light:Chemical:Mechanical | 6 | 95 | 16 | 2.489 | 0.0243 |
| Residuals | 189 | 1198 | 6 | | |

**Table S1**. Three-way ANOVA from actinula larva settlement study (7x2x2 split-plot RCBD)

| | DF | Sum Sq | Mean Sq | F-value | p-value |
|---|---|---|---|---|---|
| Block:Light | 54 | 326.9 | 6.1 | 1.1615 | 0.235 |
| Block:Chemical | 9 | 178 | 19.8 | 3.7942 | 0.0002193 |
| Block:Mechanical | 9 | 128.5 | 14.3 | 2.7402 | 0.0051625 |
| Residuals | 171 | 891.3 | 5.2 | | |

**Table S2**. Block interaction investigation from actinula larva settlement study (7x2x2 split-plot RCBD)

| | DF | Sum Sq | Mean Sq | F-value | p-value |
|---|---|---|---|---|---|
| Block | 9 | 424.0 | 47.11 | 8.942 | 4.2e-08*** |
| Light Condition | 6 | 131.2 | 21.87 | 4.151 | 0.00168** |
| Light vs Dark | 1 | 52.0 | 52.1 | 9.871 | 0.00272** |
| High vs Low | 1 | 1.1 | 1.12 | 0.213 | 0.64652 |
| G vs B&R | 1 | 57.7 | 57.69 | 10.948 | 0.00167** |
| G*Intensity | 1 | 11.0 | 11.04 | 2.095 | 0.15352 |
| B vs G&R | 1 | 2.5 | 2.49 | 0.473 | 0.4944 |
| B*Intensity | 1 | 9.7 | 9.69 | 1.839 | 0.1807 |
| R vs G&B | 1 | 36.2 | 36.19 | 6.868 | 0.01137* |
| R*Intensity | 1 | 0.0 | 0.04 | 0.008 | 0.9275 |
| Residuals | 54 | 284.5 | 5.27 | | |

**Table S3**. Photosensory contrasts in light only treatment from the actinula larva settlement study.

| | DF | Sum Sq | Mean Sq | F-value | p-value |
|---|---|---|---|---|---|
| Light:Chemical | 6 | 43 | 7 | 1.150 | 0.33379 |
| Light vs Dark | 1 | 23 | 23 | 3.6 | 0.05898 |
| High vs Low | 1 | 15 | 15 | 2.375 | 0.12463 |
| G vs B&R | 1 | 2 | 2 | 0.262 | 0.60911 |
| G*Intensity | 1 | 3 | 3 | 0.433 | 0.51130 |
| B vs G&R | 1 | 3 | 3 | 0.433 | 0.5113 |
| B*Intensity | 1 | 1 | 1 | 0.199 | 0.6557 |
| R vs G&B | 1 | 0 | 0 | 0.021 | 0.88423 |
| R*Intensity | 1 | 0 | 0 | 0.045 | 0.83283 |
| Light:Mechanical | 6 | 14 | 2 | 0.377 | 0.89337 |
| Light vs Dark | 1 | 4 | 4 | 0.682 | 0.40966 |
| High vs Low | 1 | 0 | 0 | 0.045 | 0.83283 |
| G vs B&R | 1 | 0 | 0 | 0.070 | 0.79185 |
| G*Intensity | 1 | 9 | 9 | 1.398 | 0.23815 |
| B vs G&R | 1 | 0 | 0 | 0 | 0.9985 |
| B*Intensity | 1 | 1 | 1 | 0.171 | 0.6795 |
| R vs G&B | 1 | 0 | 0 | 0.045 | 0.83283 |
| R*Intensity | 1 | 4 | 4 | 0.591 | 0.44268 |
| Light:Chemical:Mechanical | 6 | 95 | 16 | 2.514 | 0.02228* |
| Light vs Dark | 1 | 30 | 30 | 4.717 | 0.03083* |
| High vs Low | 1 | 0 | 0 | 0.021 | 0.88538 |
| G vs B&R | 1 | 31 | 31 | 5.015 | 0.02604* |
| G*Intensity | 1 | 2 | 2 | 0.346 | 0.55672 |
| B vs G&R | 1 | 3 | 3 | 0.550 | 0.4590 |
| B*Intensity | 1 | 0 | 0 | 0.052 | 0.8200 |
| R vs G&B | 1 | 56 | 56 | 8.886 | 0.00317** |
| R*Intensity | 1 | 4 | 4 | 0.666 | 0.41513 |
| Residuals | 243 | 1525 | 6 | | |

**Table S4**. Photosensory contrasts for all interactions from the actinula larva settlement study.

| Stage 5 significantly upregulated GO terms | Stage 6 significantly upregulated GO terms |
|---|---|
| GO:0005102 | GO:0005102 |
| GO:0005520 | GO:0005520 |
| GO:0005576 | GO:0005576 |
| GO:0005576 | GO:0005576 |
| GO:0005520 | GO:0009405 |
| GO:0005576 | GO:0005576 |
| GO:0005102 | GO:0005506 |
| GO:0004222 | GO:0009055 |
| GO:0008270 | GO:0020037 |
| GO:0005576 | |
| GO:0009405 | |
| GO:0005506 | |
| GO:0009055 | |
| GO:0020037 | |
| GO:0005576 | |
| GO:0006952 | |
| GO:0005102 | |
| GO:0005576 | |
| GO:0005576 | |
| GO:0009405 | |
| GO:0005506 | |
| GO:0009055 | |
| GO:0020037 | |
| GO:0005102 | |

**Table S5**. Actinula stages 5 and 6 significantly upregulated GO terms used in ReviGO analysis [1].
